# A role of hypoxia inducible factor 1 alpha in Mouse Gammaherpesvirus 68 (MHV68) lytic replication and reactivation from latency

**DOI:** 10.1101/791616

**Authors:** Darlah M. López-Rodríguez, Varvara Kirillov, Laurie T. Krug, Enrique A. Mesri, Samita Andreansky

## Abstract

The hypoxia inducible factor 1 alpha (HIFIα) protein and the hypoxic microenvironment are critical for infection and pathogenesis by the oncogenic gammaherpesviruses (γHV) such as Kaposi’ Sarcoma-associated Herpes Virus (KSHV) and Epstein-Barr virus (EBV). However, understanding the role of HIFIα during the virus life cycle and its biological relevance in the context of host pathogenesis has been challenging due to the lack of animal models for human γHV. To study the role of HIFIα we employed the murine gammaherpesvirus 68 (MHV68), a rodent pathogen that readily infects laboratory mice. We show that MHV68 infection induces HIFIα protein and HIFIα-responsive gene expression in permissive cells. Deletion of HIFIα reduces virus production due to a global downregulation of viral gene expression. Most notable was the marked decrease in many viral genes bearing hypoxia regulatory element (HRE) such as viral G-Protein Coupled Receptor (vGPCR), which is known to activate HIF1α transcriptional activity during KSHV infection. Intranasal infection of HIF1α^LoxP/LoxP^ mice with MHV68 expressing Cre-recombinase impaired virus expansion during early acute infection and affected lytic reactivation in the splenocytes explanted from mice. Moreover, low oxygen conditions accelerated lytic reactivation and enhanced virus production in MHV68 infected splenocytes. Thus, we conclude that HIFIα plays a critical role to promote virus replication. Our results highlight the importance of the mutual interactions of the oxygen-sensing machinery and gammaherpesviruses in viral replication and pathogenesis.

**AUTHOR SUMMARY:** The host oxygen sensing machinery including the HIF1α pathway is important during the viral life cycle of oncogenic gammaherpesviruses such as KSHV and EBV. However, due to the host specificity, the effects of HIF1α in herpes biology is limited to studies with *in vitro* systems. Here, we study the role of HIF1α using the mouse gammaherpesvirus 68 (MHV68) that readily infects laboratory mice. We demonstrate that MHV68 infection upregulates HIF1α during replication and inactivation of HIF1α transcriptional activity significantly decreased viral genes expression which results in impaired virus production *in vitro*. In vivo deletion of HIF1α impaired viral expansion during acute infection and affected reactivation from latency. These results show the importance of the interplay with the oxygen-sensing machinery in gammaherpesvirus infection and pathogenesis, placing the MHV68 mouse model as a unique platform to gain insight into this important aspect of oncogenic gamma-herpesviruses biology and to test HIF1α targeted therapeutics.

## INTRODUCTION

Many pathogenic viruses need to adapt to different physiological oxygen levels for efficient infection of the host by controlling the host’s oxygen sensing transcriptional machinery centered around the regulation of the hypoxia inducible factors, the main transcriptional regulators of the hypoxia-stimulated genes. Hypoxia Inducible Factor 1 alpha (HIF1α) is an eukaryotic cellular transcription factor whose main role is to support adaptation of cells and tissues to lower oxygen concentrations. Hypoxic cells react by upregulating genes to enable oxygen delivery, increase glucose uptake and anaerobic metabolism to facilitate cell and tissue survival [1,2]. HIFIα is tightly regulated by oxygen levels within the cell. In the presence of oxygen, HIFIα is rapidly targeted for degradation by ubiquitin complex via proline hydroxylation by hydroxylase dioxygenases [2]. When oxygen demand exceeds oxygen supply, HIFIα protein is no longer degraded and is translocated to the nucleus. Here, HIFIα binds the constitutively expressed HIFIβ forming a heterodimeric helix-loop-helix transcriptional complex. The HIF1 heterodimer recognizes DNA-binding motif known as hypoxic response elements (HRE) within the promoter of target genes. This leads to the expression of proteins such as vascular endothelial growth factor (VEGF), glucose transporters and erythropoietin required to adapt to low oxygen levels.

Activation of HIFIα protein has been observed during virus infection, leading to metabolic adaptation and allowing viral replication. Several viruses such as Epstein Barr Virus (EBV) [3], Human Cytomegalovirus [4], Respiratory Syncytial Virus (RSV), Varicella Zoster Virus (VZV)[5], John Cunningham Virus (JCV) [6] and Influenza A [7] are now known to upregulate HIFIα under normoxia. Particularly, oncogenic human gammaherpesviruses such as Kaposi sarcoma-associated Herpes Virus (KSHV) and Epstein Barr Virus (EBV) have evolved to exploit this component of the oxygen-sensing machinery for their survival and persistence in the host [8–13]. Kaposi’ Sarcoma (KS), an angiogenic spindle-cell sarcoma caused by KSHV, predominantly develops in lower extremities which have relatively low oxygen concentration [14–17]. KSHV infection and specific viral genes increase the levels of HIF1α and its transcriptional activity, allowing a viral-driven regulation of host processes critical for angiogenesis, glycolysis, in addition to viral gene regulation and replication [18–23]. During latency, KSHV infection imparts a hypoxic signature to infected cells [24]. *In vitro* experiments have demonstrated that HIF1α plays an important role in lytic reactivation of KSHV and EBV from latently infected cell lines by binding to the promoter of the immediate early viral gene RTA or Replication and Transcription Activator [11,12,25,26]. Similarly, exposure of latently infected mouse B-cell lymphomas with mouse gammaherpesvirus 68 to hypoxia conditions triggered lytic reactivation of MHV68-increased transcription activity of RTA [27]. Furthermore, Latency Associated Nuclear Antigen (LANA), a key viral protein, enhances HIF1α transcription and cooperates with RTA to promote lytic replication [8].

Infection with herpesviruses leads to lytic replication followed by latency establishment in the host. Viral latency in infected cells sustains the persistence of the virus during its lifetime, while lytic replication from latently infected cells permits the spread of the virus. Given the host-specific nature of human gammaherpesviruses, the role of HIF1α in pathogenesis is difficult to elucidate as they exhibit limited lytic replication *in vitro* and there is no established small animal model of infection. Opportunely, Murine gammaherpesvirus 68 (MHV68) undergoes lytic infection upon *de novo* infection in permissive cells and readily infects laboratory mice. MHV68 is genetically related to KSHV and encodes many homologues of KSHV that are required for both lytic and latent stages of the virus life cycle [28]. Thus, our objective was to elucidate the role of HIF1α during MHV68 virus life cycle using both *in vitro* and *in vivo* infection models.

We report that MHV68 infection of permissive cells upregulated HIF1α transcription and led to upregulation of its protein levels. Genetic ablation of HIF1α transcription activity decreased production of mature virions and expression of several viral HRE-containing viral genes. Ablation of HIF1α transcription activity *in vivo* by intranasal infection of HIF1α^LoxP/LoxP^ mice with an MHV68 virus expressing Cre-recombinase impaired virus expansion in lungs early in infection and affected reactivation of the virus during latency. These findings establish the role of HIF1α during gammaherpesvirus pathogenesis in an inherent host.

## RESULTS

### MHV68 infection upregulates HIFIα expression and transcriptional activity

We first determined whether MHV68 upregulates HIF1α during virus infection in culture. The mouse fibroblast cell line NIH 3T12 was infected with a wild type MHV68 strain in normoxia (21% O_2_) and HIF1α mRNA and protein levels were analyzed by western blot and qRT-PCR (Figure 1). Fig 1 shows that MHV68 infection upregulates HIF1α protein which increases over time. Cobalt chloride (CoCl_2_), a hypoxia mimic was used as positive control [29]. Accumulation of HIF1α protein correlated to a 6-fold increase in HIF1α mRNA levels (Figure 1B) at 24 hpi when compared to uninfected cells indicating that induction of HIF1α activity by MHV68 occurs together with activation of transcription. Moreover, transcription of HIF1α was dependent on viral gene expression, as we did not detect HIF1α mRNA upregulation when cells were exposed to UV-inactivated virus (Figure 1B). We next sought to determine whether upregulation of HIF1α during MHV68 infection activates HIF1α mediated transcription of host HIF1α regulated genes, which containing HRE-binding sites at the regulatory region using an HRE-dependent luciferase reporter and a dual-luciferase assay. 3T12 cells were transfected with reporter or control vectors and then infected under normoxia (21% O_2_) and hypoxia (1%O2) at different MOI. We found an increase in firefly luciferase reporter activity 24 hpi in cells infected with MHV68 in comparison to uninfected controls within 24 hpi (Figure 1C) which was higher in hypoxic than normoxic conditions suggesting that both infection and hypoxic conditions contribute to enhancement of HIF1α transcription activity. Upregulation of HIF1α by oncogenic gammaherpesviruses such as KSHV and EBV is central to the induction of metabolic reprogramming which occurs via upregulation of HIF1α regulated genes such as glucose transporter 1 (GLUT-1), glucose-6 phosphate isomerase (GPI) and pyruvate kinase (PKM). These are key enzymes required for energy production for cellular adaptation to oxygen. We therefore determined if HIF upregulation by MHV68 lead to increase in transcription of these metabolic HIF-target genes using qRT-PCR. Transcription of genes was increased 7 to 5-fold in MHV68 infected cells (Figure 1D) and was dependent on virus infection as cells exposed to UV-irradiated virus failed to induce upregulation of HIF1α-regulated genes. Taken together, the data depicted in Figure 1 shows that MHV68 infection upregulates HIF1α levels and transcriptional activity.

**FIG 1.**
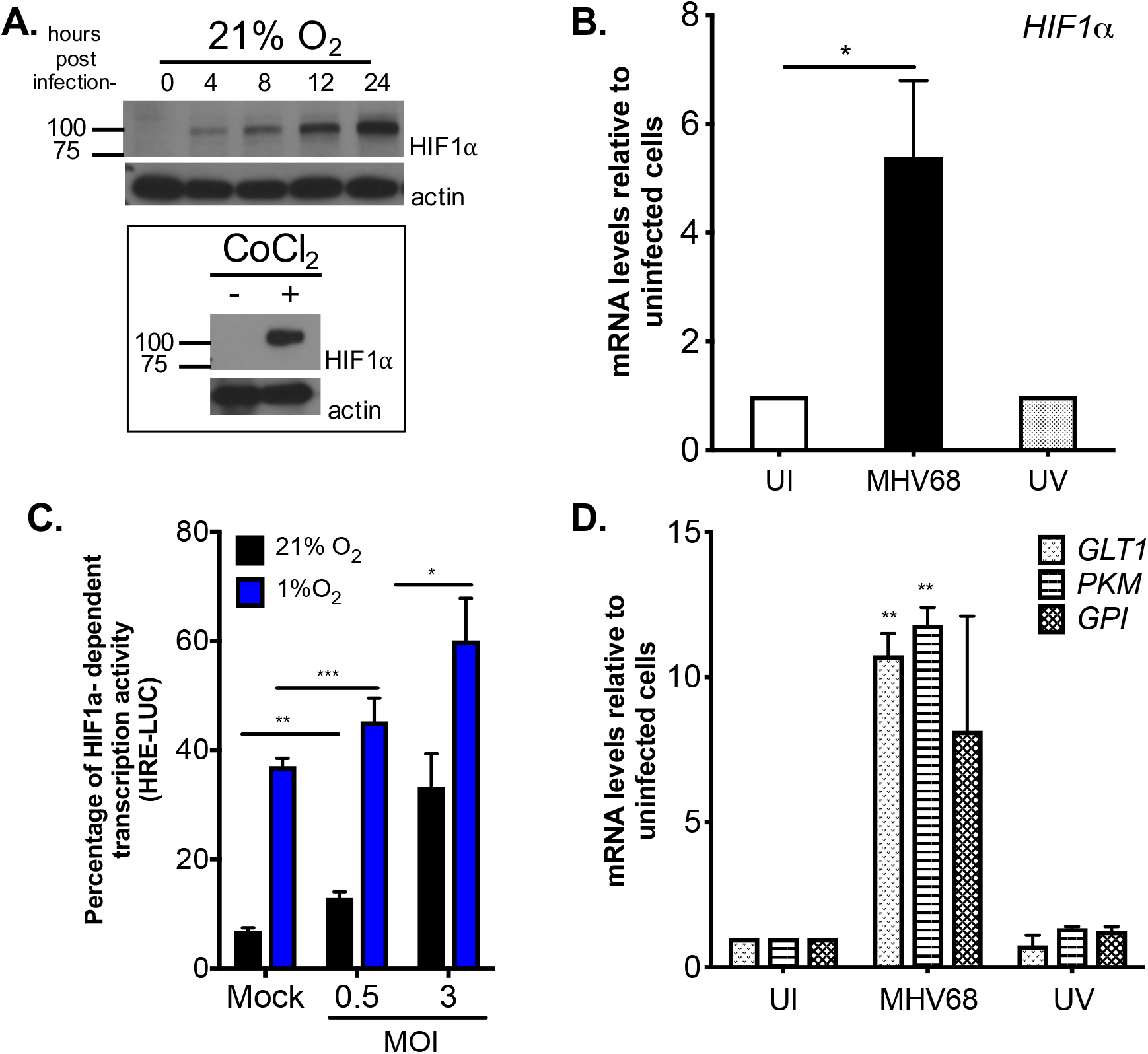
MHV68 infection upregulates expression of HIF1 alpha. (**A**) 3T12 fibroblasts were infected with a wild type strain of MHV68 (WUMS) (5 MOI) at 21% O_2_ (cell culture incubator) and 30ug of protein lysates were analyzed by western blot for the expression of HIF1α protein at different time-points. A second set of cells were treated for 8 hours with the hypoxia-mimic CoCl_2_, which served as positive control (enclosed right panel). (**B**) HIF1α mRNA at 24 hpi expressed as fold-change in cells infected with MHV-68 or UV-irradiated virus relative to uninfected cells. Graph depicts results of one out of three experiments performed in triplicates. Statistical analysis by Student’s t-test. *, *p<0.05*. (**C**) 3T12 cells were transiently transfected with pGL2 vector (0.5 ug) with dual reporter system (Renilla, 0.5 ug) which contains the three hypoxia response elements from the *Pgk-1gene* for 12 hours followed by MHV68 infection (MOI = 0.5 and 3.0). Cells were transferred to, 21% O_2_ or normoxia (black bar) or 1% O_2_ or hypoxia (blue bar) and HRE-driven luciferase activity was measured at 24 hpi. Statistical analysis by Multiple Student’s t-test of one experiment performed in triplicate. Graph is representative out of three experiments. *, *p<0.05*. **, *p<0.01*. ***, *p<0.005* (**D**) mRNA levels of HIF1 alpha targeted host genes such as *GLT1, PKM* and *GPI* were measured by qPCR at 24 hpi. Uninfected and UV-irradiated MHV68 virus were used as negative controls. *GLT1=* glucose transporter 1, *PKM=* pyruvate kinase, *GPI*= glucose-6-phosphate isomerase. Graph represents one experiment out of three. Statistical significance of differences in fold-change was determined by Multiple Student’s t-test of triplicates. **, *p<0.01*.

### Genetic Ablation of HIF1α DNA binding domain suppresses hypoxia driven transcription

Upregulation of HIF1α protein during MHV68 infection indicates that this transcription factor plays a role during virus replication. We therefore sought to evaluate the impact of HIF1α on lytic replication and viral expression in knock-out cells. We obtained primary MEFs from transgenic knock-in mouse (B6.129-Hif1*α^tm3Rsjo^*/J), with exon 2 of the *HIFIα* gene flanked by 34bp specific *LoxP* sites (HIF1αLoxP MEFs) [30]. Exon 2 encodes the DNA binding region required for the dimerization of the protein in the nucleus and transcription of HIF1 target genes. A cre-recombinase expressing lentivirus was employed to transduce HIF1αLoxP MEFs followed by selection for resistance to the antibiotic Blasticidin. We first characterized both HIF1α wild-type (WT= MEFs from HIF1αLoxP transgenic mice) and HIF1α Null cells (Null= HIF1αLoxP MEFs expressing Cre-recombinase) by performing mRNA analysis with specific HIF1α oligonucleotides for HIF1α cDNA. Exon 2 deletion was detected as a fragment size shift to 400bp in HIFa Null cells in contrast to the complete 600bp PCR product spanning exon 1 to exon 5 in non-transduced HIF1αLoxP MEFs (Figure 2B). Also, no amplification of exon 2 was detected by qPCR in Null cells when compared to WT MEFs and no change of expression was observed in Exon 4/5 transcripts (Figure 2B).

**FIG 2.**
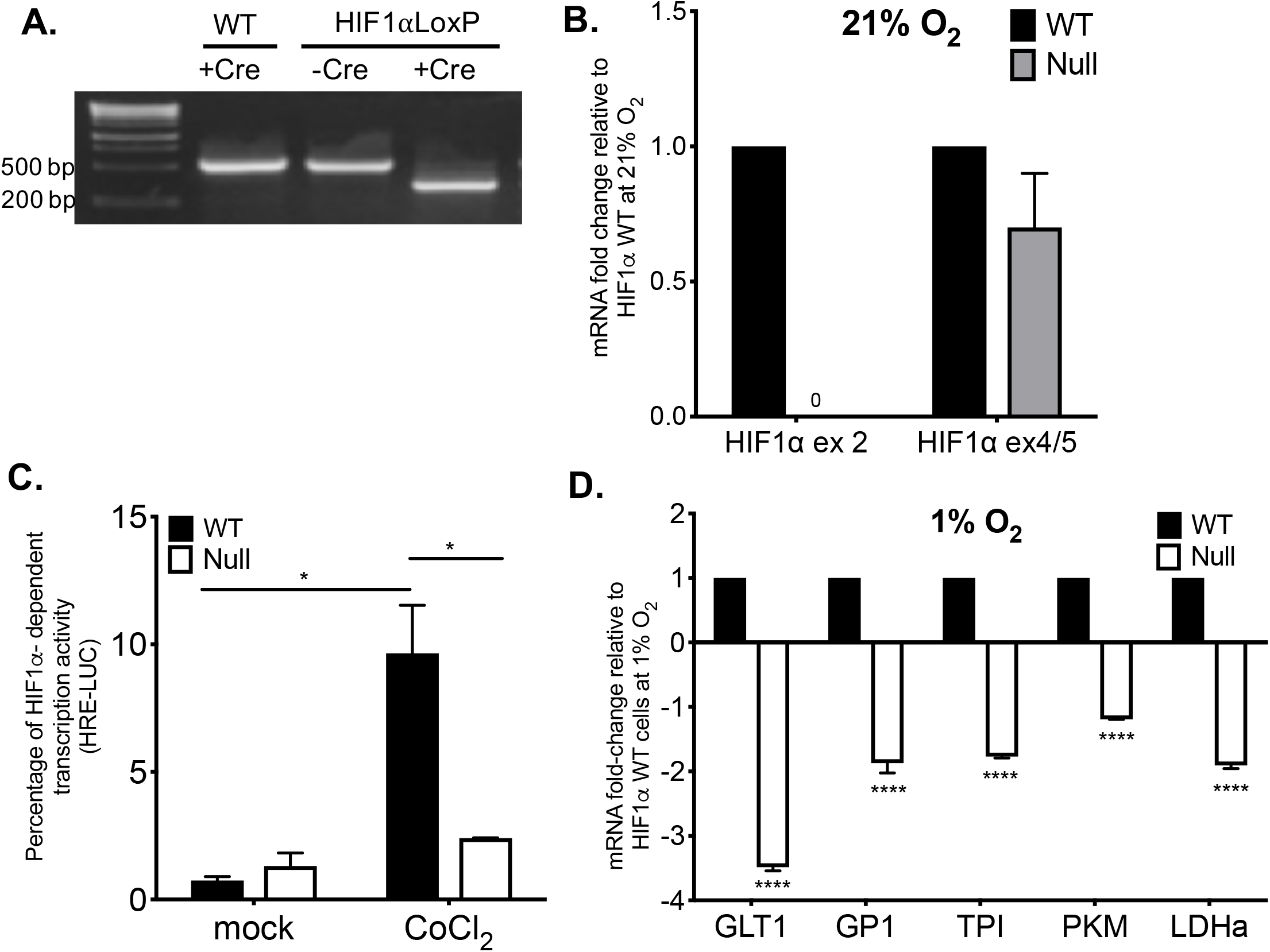
Deletion of HIF1α DNA-binding domain suppresses HRE-dependent transcription in hypoxia. (**A-D**) Mouse embryonic fibroblasts (MEFs) were isolated from 13.5-day old embryo from B6.129-*Hif1a^tm3Rsjo^*/J (HIF1αLoxP) and C57BL/6J (WT) and were immortalized by culturing cells over 30-35 generations. Immortalized HIF1αLoxP MEFs cells were transduced with a lentivirus vector expressing Cre-recombinase (Lenti-Cre) and selected with Blasticidin. MEFs (WT) isolated from parental mice was used as corresponding control for all experiments. (**A**) Excision of exon 2 was detected by amplification of gene fragment spanning exon 1 to exon 5 by PCR. A 400 bp fragment corresponds to the excised exon 2 in Null MEFs (+CRE) in comparison to 600 bp fragment (-CRE) in HIF1αLoxP MEFs. (**B**) HIF1α mRNA expression in WT and Null cells were measured by qPCR with primers from Exon 2 region. Exon 4/5 from HIF1α primer was used as corresponding control and was detected in both WT and Null cells. (**C**) WT and Null MEFs were either treatment with the hypoxic mimic cobalt chloride (CoCl_2_) to induce HRE-driven luciferase expression for 8 hours or left untreated. Graph represents data from one experiment performed in triplicates. *, p<0.05 by Student’s t–test. (**D**) WT and Null MEFs were exposed to 1% O_2_ and HIF1 alpha target genes such as Glutamate transporter (*GLT*), Glucose-6-Phosphate Isomerase (*GPI*), Triose-phosphate Isomerase (TPI), Lactate Dehydrogenase A (*LDHa*), Pyruvate Kinase M1/2 (*PKM*) were measured by qPCR. Data was normalized as relative fold-change in WT. Statistical significance was determined in GraphPad prim by Unpaired Student t-test of one experiment performed in triplicates. ****, p<0.0001. Graph represents data from one experiment performed in triplicates.

Null cells were further analyzed to confirm that they lacked HIF1α transcriptional activity using an HRE-luc reporter and qRT-PCR for HIF1α-regulated genes as done in Fig 1C and 1D. Luciferase signal was 10-fold less in Null cells following 8-hour treatment with the hypoxia mimic CoCl_2_, indicating HIF1α dependent activity was impaired. In addition, transcription of HIF1α-regulated glycolytic genes were also verified by qRT-PCR in Null cells. Each HIF1α transcript exhibited a significant decrease in expression after 8-hour treatment in 1% O_2_ conditions, confirming that HIF1α protein was inactive in Null cells.

### Absence of HIF1α activity impairs MHV68 replication *in vitro*

Next, we assessed whether absence of HIF1α transcription activity plays a role during the virus lytic replication. HIF1α wild-type (WT) and Null MEFs were infected with high and low MOI of MHV68 virus, viral supernatants were harvested at different times post infection (dpi) and infectious virus was assayed by plaque assay. Comparing virus titers in WT and Null cells inoculated with varying MOI, virus production was decreased uniformly in the absence of HIF1α. The data is represented as mean plaque forming units per ml (Figure 3). As shown in Figure 3A, time-course infection of Null cells at 5.0 MOI showed a slight reduction while a lower infection of 0.5 MOI had a significant decreased in virus production at later time-points. These results indicate that cells lacking HIF1α significantly reduce kinetics of viral replication and the extent of viral release or viral burst size in both MEFs is dependent on the input virus (Figure 3A). Thus HIF1α play a significant role for efficient production of infectious particles during MHV68 replication.

**FIG 3.**
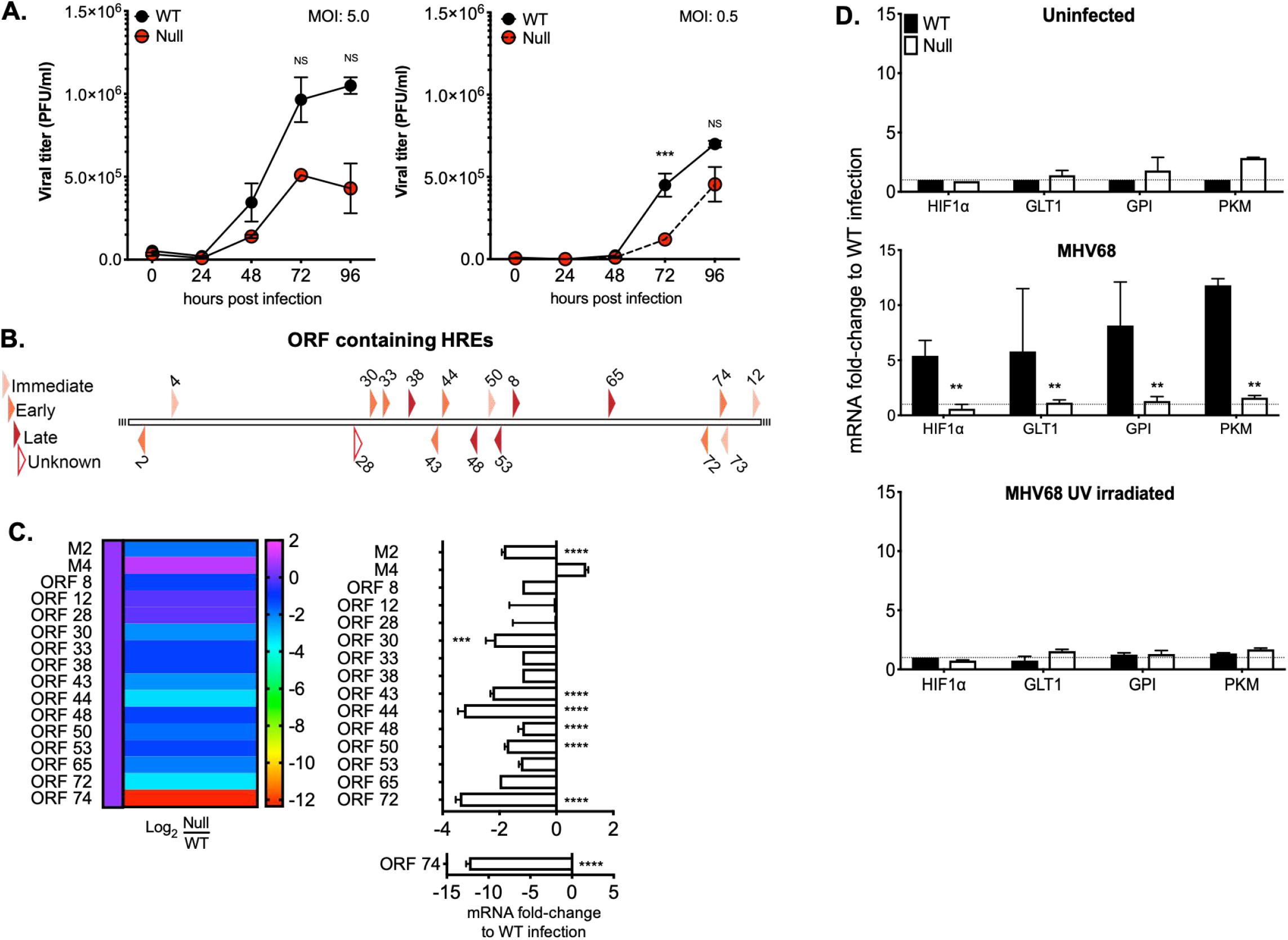
Viral replication is compromised in the absence of HIF1α and is required for transcriptional activity of HRE-containing viral and host genes. (**A**) Stably transduced WT and Null MEFs were infected with wild type MHV68, WUMS strain at 5 MOI (**Left**) and 0.5 MOI (**Right**) in normoxia. Virus supernatants were collected at 0, 24, 48, 72 and 96 hpi and assayed for released virus by plaque assay on 3T12 cells. Graph represents one of three independent experiments performed in triplicates. Statistical significance was determined in Graph Pad Prism by multiple Student’s t-test. ***, *p< 0.005*. NS abbreviates no statistical significance. (**B**) Schematic diagram of MHV68 open reading frame promoters containing 1 or 2 potential HIF1α binding sites (R-CGTG) was analyzed using TRANSFAC database. The diagram categorizes time of expression upon *de novo* lytic infection of MHV68 in 3T12. White polygon represents an unknown function. (**C**) WT and Null MEFs were infected with wild type MHV68 (MOI 5.0) and incubated at 21% O_2_. RNA was isolated 24 hpi and changes in viral open reading frames (ORF) was measured by qPCR. ΔΔCt was expressed as fold change and normalized against WT MEFs infection. Data is representative of one out of three experiments. Statistical significance determined by multiple t-test using the Holm-Sidak method, with alpha= 0.05 on one experiment performed in triplicates. (**D**) WT and Null MEFs were infected with wild type MHV68 as in 4B. RNA was isolated 24 hpi and levels of mRNA for *GLT1* (glucose transporter 1), *PKM* (pyruvate kinase) and *GPI* (glucose-6-phosphate isomerase) were determined by qPCR; ΔΔCt was expressed as fold change and normalized against uninfected HIF1α WT MEFs. Data is representative of one out of three experiments. Unpaired t-test with Welch’s correction *P*< 0.01 on one experiment performed in triplicates.

### Absence of HIF1α impairs viral gene expression in MHV68

The murine gammaherpesvirus shares significant homology with human gammaherpesviruses such as KSHV and EBV and encodes for several viral orthologs. KSHV genes containing the HRE consensus sites (5’-ACGTG-3’) are transcriptionally upregulated by HIF1α in the presence of hypoxia [31] and HIF1α regulates viral persistence by binding HRE homologs located throughout the genome [25]. Thus, we analyzed MHV68 viral promoters containing the consensus HIF1α binding motif with Biobase TRANSFAC database. The transcription element search system was employed to identify potential transcription binding sites containing the string site RCGTG within 500 base pairs upstream of the starting codon of all the MHV68 open reading frames. The results, depicted as a diagram in Figure 3B, predicted 17 viral promoters with HREs in all classes of MHV68 genes (immediate early, early and late), including the KSHV orthologs of open reading frames (ORFs) 43, 44, 50, 73 (LANA) and 74 (vGPCR) which belong to the hypoxia responsive KSHV clusters [25].

A qRT-PCR was performed in infected WT and HIF1α Null cells to measure mRNA levels of these 17 MHV68-HRE containing and non-containing ORFs. Viral mRNA was harvested from infected cell lysates 24 hpi since low cytolysis is observed and cell associated viral RNA is still intact at this time. Absence of HIF1α activity decreased transcription of many HRE and non-HRE containing viral genes in Null cells when compared to the transcript levels of WT MEFs (Figure 3C and Supplementary Figure 1). Within the HRE-containing viral genes, the most notable downregulation was observed for the viral G protein-coupled receptor (vGPCR/ORF74), a KSHV viral gene known to regulate HIF1α transcriptional activity and angiogenesis in KS [21–23]. In addition, ORF72, viral cyclin D homolog and ORF73, latency associated nuclear antigen LANA were reduced 3-fold Several HRE-containing viral genes designated for viral replication such as ORF44, a component of DNA helicase-primase complex and ORF65, a DNA packaging protein were downregulated 2-fold and was statistically significant. Taken together our results indicate that virally-induced HIF1α participates in the expression of many HRE-containing and non-containing promoters regulating early and late genes necessary for optimal growth kinetics during MHV68 replication.

### HIF1α activity is required to induce host genes that affects MHV68 replication

MHV68 infection of 3T12 cells increased transcription of glycolytic genes (Figure 1D). This data is in line with observation that herpes virus infections induce glycolysis through anabolic pathway in order to support increased demand on cellular translation machinery required during viral replication and also to maintain latently infected cells [32].

We undertook analysis of genes involved in glucose uptake after virus infection in WT and Null cells. UV-irradiated virus and mock-infected MEFs were used for negative controls. Lytic infection upregulated up to 5-10 folds several enzymes such as glucose transporter 1 (GLT1) which is involved in glucose uptake [33], glucose-6 phosphate isomerase (GPI), which is the first enzyme in glycolytic pathway, and pyruvate kinase (PKM), which catalyzes the final step of glycolysis a wild-type infection of WT MEFs in comparison to uninfected (Figure 3D). HIF1α transcript levels were assessed as internal control during virus replication. The increase in gene expression in WT MEFs was dependent on replication of the virus as UV-inactivated virus did not induce aerobic glycolysis in WT MEFs. In contrast, glycolytic gene expression was consistently similar to uninfected in Null cells. (Figure 3D), suggesting that transcriptional activity of HIF1α is required for the induction of glycolytic enzymes during MHV68 lytic replication.

### HIF1α is necessary for optimal MHV68 replication in lower, physiological, oxygen levels

Our data demonstrates that absence of HIF1α affects viral gene expression and virion production during lytic replication in normoxia. However, oxygen levels may play a profound role during *in vivo* infection as tissues and organs are usually characterized by their unique oxygenation status. During low oxygen availability, HIF1α accumulates binding to promoter regions carrying specific HRE elements. Therefore, we speculated that lower oxygen levels along with the effects in lack of HIF1α will be more profound.

We performed a western blot analysis to determine the expression of HIF1α protein after virus infection in 3% O_2_ conditions hypoxia, since physiological levels of oxygen in many tissues ranges 3-7% [34]. 3T12 cells were infected with wild type, MHV68 in normoxia for 2 hours in 21% O_2_ and then moved to hypoxia chamber. Cell lysates were harvested 4-24hpi and HIF1α expression was analyzed by western blot analysis. In figure 4A, we show that low levels of O_2_ enhanced accumulation of HIF1α protein during virus infection.

**FIG 4.**
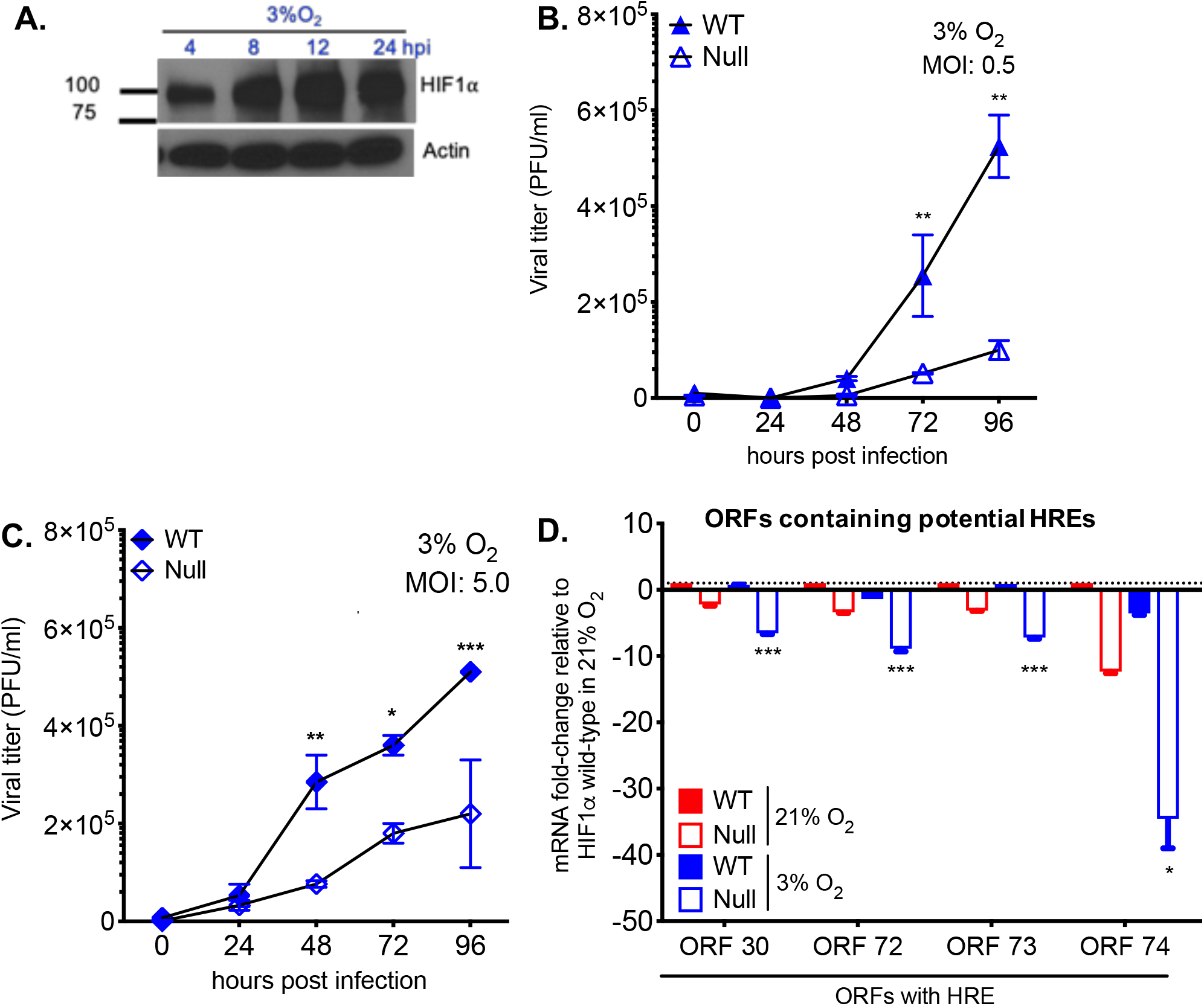
Absence of HIF1α impairs gammaherpesvirus lytic replication in low oxygen concentration. (**A**) HIF1α expression during MHV68 time-course infection (4,8,12 and 24 hpi) at 3% O_2_ was measured by western blot. (**B**) HIF1α WT and HIF1α Null MEFs were infected with MHV68 (**B**: MOI 5.0) in a single-step and (**C**: MOI 0.5) in a multi-step infection and transferred to 3% oxygen. Released virus in the supernatant was measured by plaque assay. Graph represents one of three independent experiments. Statistical significance was determined in Graph Pad Prism by Student’s t-test with n=3. **, *P*< 0.01; ***, *P*< 0.005. (**D**) Selected viral genes with statistical significance *p<0.05* of HIF1α Null 3% O_2_ normalized to HIF1α Null 21% O_2_.

The role of HIF1α on MHV68 replication at different oxygen levels was assessed in WT and Null MEFs infected with high and low MOI of virus by quantifying virion production at various times. Figure 4B demonstrates that viral expansion in the absence of HIF1α decreases especially as low MOI infection progresses under low oxygen tension with 2.3 and 4.5-fold change at 72 and 96 hpi, respectively.

To understand how oxygen level may affect the ability of HIF1α to regulate viral gene expression during virus infection, transcription analysis of HRE containing viral genes was performed 24 hpi as in Figure 4B. The data is represented by relative fold-change values which were normalized against infected WT and Null MEFs under normoxic conditions (Figure 4D). Absence of HIF1α at low oxygen levels had 10-fold reduction in expression of several HRE-containing genes such as cyclin D, LANA and vGPCR. This decrease was most notable in some HRE containing viral genes including vGPCR which was reduced 34.6-fold in Null cells when compared to WT MEFs under 3% oxygen. Non-HRE viral genes including ORF9 and ORF25 were also strongly downregulated at 3% O_2_ in the absence of HIF1α (Fig 4C and Supplementary Figure 1). Mainly, viral proteins related to viral and DNA replication (ORF9, RTA), assembly and latency associated genes such as LANA (ORF73), cyclin D (ORF72) and M2 (Supplementary Table 1) were impacted by low oxygen level conditions in Null cells.

### The role of HIF1α in MHV68 *in vivo* pathogenesis

Our data showed that HIF1α protein plays a significant role in the replication of MHV68. However, it is unknown the exact role that the HIF1α pathway plays in gammaherpesvirus pathogenesis. Since infection of mice with MHV68 provides a tractable animal model that manifests the fundamental strategies for gammaherpesvirus pathogenesis [28], we took advantage of genetic ablation of HIF1α in the HIF1α^LoxP/LoxP^ mice by infection (Figure 5A) with a recombinant MHV68 virus encoding the Cre-recombinase protein under CMV promoter (MHV68-Cre) [35]. Homozygous deletion of HIF1α is lethal for development through embryogenesis [36], we utilized Cre-LoxP strategy to generate HIF1α deletion during MHV68-Cre infection. Several studies have reported the use of engineered MHV68 encoding Cre-recombinase gene to study virus-host interaction [37–39]. Our objective was to achieve deletion of exon 2 in *HIFIα* locus in tissues by infection with a MHV68-Cre virus.

**FIG 5.**
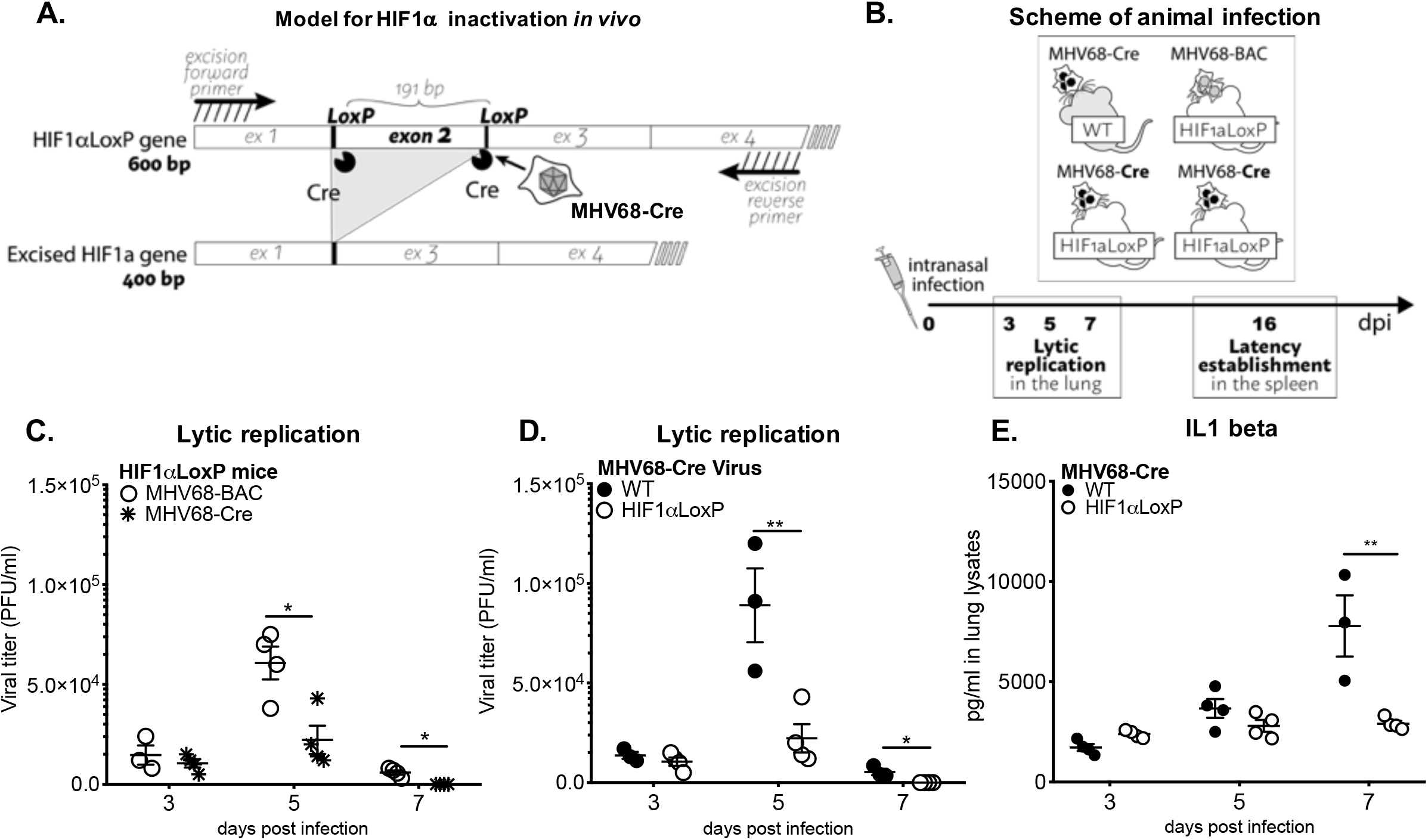
HIF1α deletion affects ***in vivo*** virus growth expansion during acute infection. (**A**) Diagrams depict HIF1α inactivation by MHV68-Cre virus infected cells. (**B**) Scheme of animal infection. (**C-D**) C57BL/6J (WT/B6) mice (n=3-4) or B6.129-*Hif1a^tm3Rsjo^/J* (HIF1αLoxP /B6) mice (n=3-4) were infected with MHV68-Cre virus by intranasal infection. Viral replication was measured from whole lung homogenates at 3, 5- and 7-days infection by plaque assay. Viral plaques were counted 5 dpi and expressed as PFU/ml. HIF1αLoxP mice were infected either with wild type MHV68 virus, (**C**) BAC-derived (n=3-4) or (**D**) MHV68-Cre virus (n=3-4) and lungs were assayed for viral titers as in 5A. Virus replication was significantly decreased on day 5 (*P<0.012*) or day 7 (*P<0.0016)*. (**E**) Both WT and HIF1αLoxP mice were infected with MHV68-Cre virus and lungs were collected as described in 5A. ELISA was performed from lung homogenates to measure IL1 beta, TNF alpha, IL6 and IFN gamma. There were no differences in cytokine production between the two mice background, except for the marked decrease in IL-1beta from lungs of HIF1αLoxP mice on day 7 (p=0.12808) in comparison to WT. Graphs represent one experiment of three independent experiments. Statistical analysis was performed in Graph Pad Prism by Multiple Student’s-t-test.

MHV68 undergoes a period of lytic replicative expansion in the respiratory tract and to a lesser extent in the spleen after intranasal infection of laboratory mice. Robust viral replication in lungs is characterized by infectious virion production and is cleared within 10-15 dpi [28]. In order to define whether HIF1α plays a role during MHV68 infection, we examined virus replication in lungs and latent virus establishment and reactivation from splenocytes. C57BL/6 WT (wild-type) and HIF1α^LoxP/LoxP^ mice were infected intranasally with 3 × 10^4^ PFU of MHV68-Cre (Figure 5B) virus. A second set of experiments were performed in HIF1α^LoxP/LoxP^ mice infected with MHV68-BAC (parental strain for the recombinant virus) to validate that transgenic mice equally supports MHV68 replication (Figure 5B).

Lungs were harvested on days 3, 5 and 7 post infection and viral titers were measured from lung homogenates. MHV68-Cre virus established infection in both WT (1.3 × 10^4^ PFU/ml ± 1.8 × 10^3^) and HIF1α^LoxP/LoxP^ mice (7.1 × 10^3^ PFU/ml ± 2.4 × 10^3^) by day 3 post-infection (Figure 5C). However, there was four-fold reduction of virus titer (2.2 × 10^4^ PFU/ml ± 7 × 10^3^) in HIF1α^LoxP/LoxP^ mice on 5 dpi when compared virus titers in C57BL/6 WT (8.9 × 10^4^ PFU/ml ± 1.9 × 10^4^) mice. The decline in viral titers continued until day 7 post infection with titer below limit of detection for HIF1αLoxP infection when compared to 5.4 × 10^3^ PFU/ml ± 1.6 × 10^3^ virus in WT mice (Figure 5C). The decrease in acute viral replication was related specifically to deletion of HIF1α activity, as viral kinetics and production were not affected in HIF1α^LoxP/LoxP^ mice infected with MHV68-BAC (wild type) virus (Figure 5C). The mean PFU/ml was 1.5 × 10^4^ PFU/ml ± 4.8 × 10^3^ on 3 dpi and 6.0 × 10^4^ PFU/ml ± 8.2 × 10^3^ on 5 dpi in these mice (Figure 5C) and was similar to viral titers observed in WT (C57Bl/6J) mice infected with MHV68-Cre virus.

Early innate immune responses to MHV68 infection is accompanied by infllammation [40–43]. Inflammatory cytokines involved in this process, include interleukin-1 (IL-1) and TNFa. Several cytokines such as IL1β, IFNβ, IL-6, TNFa and IFNγ were analyzed from lung homogenates by ELISA from WT and HIF1α^LoxP/LoxP^ mice infected with MHV68-Cre virus. Although there was a trend in reduction of cytokine production in lungs from infected HIF1α^LoxP/LoxP^ mice on day 7 when compared to C57Bl/6J mice, the levels were not statistically significant (data not shown) except for IL1β which was reduced 3.5-fold in floxed mice (Figure 5E). The reduction in proinflammatory cytokines on day 7 post infection in the absence of HIF1α activity may be due to early viral clearance reflected titers at day 5. We conclude that inhibition of HIF1α activity during acute MHV68 infection impairs virus expansion in the initial days of infection.

Similar to human gammaherpesviruses, following virus clearance in the lungs, MHV68 establishes life-long latency in the host [44,45], and the spleen is the major site of latent reservoir. The establishment of latency is observed as early as day 16 post infection, where a substantial number of splenocytes (mostly naïve B cells) can be reactivated to produce lytic virus when co-cultured *in vitro* with permissive cells [46,47]. Therefore, we determined whether HIF1α plays a role during viral latency establishment *in vivo* and reactivation *ex vivo*. C57BL/6 (WT) and HIF1α^LoxP/LoxP^ mice were infected with MHV68-Cre virus and splenocytes were harvested on days 16. The frequency of splenocytes harboring viral DNA (establishment) was determined by nested PCR. This assay has single-copy sensitivity for ORF50, which equates to one viral genome-positive cell. On the y-axis, the percentage of reaction positive for viral DNA at each cell dilution on x-axis. There were no significant differences in latency establishment in infected HIF1α^LoxP/LoxP^ (1 in 2,880 cells) and WT mice (1 in 2,346 cells) on 16 dpi (Figure 6A). The same splenocytes were assayed to measure the frequency of reactivating virus by *ex vivo* limiting dilution assay (LDA). Splenocytes were diluted 10-fold and co-cultured with primary MEFs for two weeks. The number for the frequency of cells reactivating was determine based on the Poisson distribution which predicts that 0.1 PFU per well should result at 63% percent reactivation of wells positive for cytopathic effect (CPE) [35].

**FIG 6.**
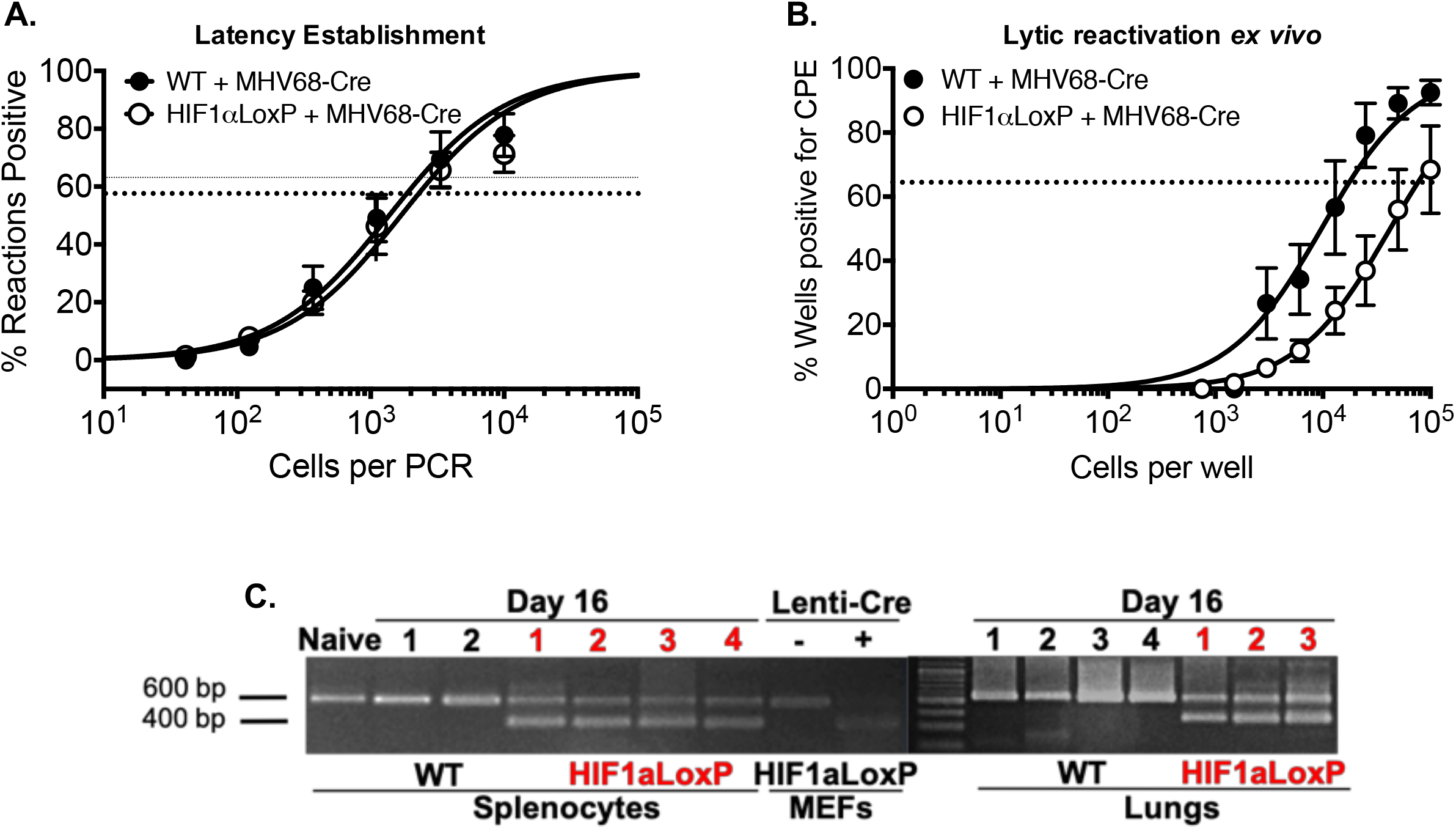
Excision of HIF1α decreases viral reactivation from latency in infected splenocytes in vivo. C57BL/6J mice (n=3-4) or B6.129-*Hif1α^tm3Rsjo^*/J (n=3-4) were infected with MHV68-Cre virus by intranasal infection. Splenocytes were harvested on day 16. (**A**) Limiting-dilution PCR was performed from infected WT and HIF1αLoxP splenocytes with two rounds of PCR were performed against MHV68 ORF50. There were no statistical differences (*P=0.4350*) between the two groups of mice infected with MHV68-Cre virus. (**B**) *Ex vivo* reactivation by limiting-dilution assay was performed to determine the frequency of infected WT and HIF1αLoxP splenocytes that harbor the viral genome. The frequency of cells reactivating the virus in HIF1αLoxP mice were less (1 in 16,818 splenocytes), when compared to C57BI/6J mice (1 in 73,181 splenocytes) and was statistically significant (*P= 0.0287)*. For both limiting-dilution assays, curve fit lines were derived from nonlinear regression analysis. Symbols represent the mean percentage of wells positive for virus CPE +/− the standard error of the mean. (The dotted line represents 63.2%, from which the frequency of cells reactivating virus was calculated based on the Poisson distribution. Statistical significance was determined in Graph Pad Prism by Student’s t-test. *, *p*< 0.05. (**C**) RNA from 10^7^ splenocytes (25ng per PCR rxn) and Lung homogenates (25ng per PCR rxn) of WT and HIF1αLoxP mice were analyzed for excision of HIF1α exon 2 by PCR and the products were run on DNA agarose gels. Tissue from an uninfected naïve HIF1αLoxP mouse was used as negative control. A 400bp fragment was observed only HIF1αLoxp and not in parental WT mice when infected with MHV68-Cre virus. It is important to note that both lungs and splenocytes are major sites of MHV68 replication.

In contrast to latency establishement, significantly less splenocytes reactivated in HIF1αLoxP (1 in 73,181 splenocytes) mice when compared to WT infection (1 in 16,818, *P= 0.0287*) upon *ex vivo* culture as shown in Figure 6B. We also confirmed that transgenic background did not affect the frequency of viral reactivation by LDA assay from splenocytes harvested from HIF1α^LoxP/LoxP^ mice infected with wild type (MHV69-BAC) virus. We validated the excision of exon 2 *in vivo* on RNA isolated from lung tissue and bulk splenocytes on16 dpi. A 400 bp PCR product corresponding to excised HIF1α gene was observed only in lungs and splenocytes of HIF1α^LoxP/LoxP^ mice infected with MHV68-Cre virus (Figure 6C). A 600 bp product relating to full-length HIF1α gene in uninfected (naïve) or in WT mice infected with the same virus confirmed that cre-recombinase was functional *in vivo*.

### *Ex vivo* reactivation of MHV68 infected splenocytes in hypoxia enhances virus production

We found that reactivation frequency was affected by virus-specific inactivation of HIF1α. These results suggest that HIF1α plays a role during the latent to lytic switch *ex vivo*. This is consistent with our results in Figure 3 showing that HIF1α deletion impairs lytic replication and studies showing that hypoxia conditions induced expression of lytic antigens in MHV68 latently infected cells [27]. However, the role of the hypoxia/HIF pathway during viral reactivation of gHV latently infected cells following a primary infection in a natural host has not been explored. We first assessed whether low oxygen culture conditions could affect the frequency of reactivating MHV68 positive cells following latency establishment of WT mice and wild-type MHV68 *in vivo*. The limiliting dilution assay was carried out at 3% O_2_ and compared to 21% O_2_ after 7-days in culture as in Figure 6 (See Materials and Methods). There was no difference in the frequency of reactivation, 1 in 42,743 splenocytes reactivated in normoxia while low oxygen yields 1 in 29,498 splenocytes (Fig. 7A). This indicates that MHV68 positive splenocyes switch to the lytic phase at a similar rate regardless of oxygen concentration. We then examined whether low oxygen levels would increase the amount of viral particles released during reactivation of cells latently infected with MHV68. Splenocytes from WT mice 16 dpi were plated at different ratios on to 2 × 10^5^ MEFs then incubated at 21% O_2_ or 3% O_2_. The released infectious virus in supernatants was titered by plaque assay. On day 4, no infectious particles were detect in co-cultures from normoxic conditions (21%O_2_) regardless of splenocytes-to-MEFs ratio and within the limits of detection by the assay. In contrast, as shown in Figure 7B, infectious virus was already detectable on all splenocytes-to-MEFs cultured in low oxygen conditions. Virus production from 10^4^ reactivating splenocytes to 10^5^ MEFs (Figure 7B-Left) was not detected in 21% O_2_ cultures. However, at 3% O_2_ concentration, virus prodution continued to produce on average 1.3 × 10^3^ PFU per ml ± 187.5 on day 4, 8.3 × 10^3^ PFU per ml ± 7.4 × 10^3^ on day 5 and 2.6 × 10^5^ PFU per ml ± 1.3 × 10^3^ on day 6. Likewise, as shown in Figure 7B (Right), 10^6^ reactivating splenocytes (10:1) had the highest increment in 1% O_2_ conditions of up to 500-fold change on day 5 and 10^5^ reactivating-splenocytes (1:1) with 100-fold boost. No virus production was detected following 3 days in co-culture supernatant from either condition but mRNA analysis by qPCR revealed significantly higher RTA expression in hypoxic condition was when normalized to normoxic reactivation regardless of splenocytes to MEFs ration (Fig 7C). Thus, our data shows that low oxygen provide cellular conditions that accelerate and enhance viral production during reactivation from latency in the B-cell lineage the primary reservoir of gammaherpesviruses. This is aligned with our finding in Figure 6B showing that reactivation is impaires by loss of HIF1α further reinforcing the idea that hypoxia and the HIF1α pathway plays a role in gammaherpesvirus reactivation from latency.

**FIG 7.**
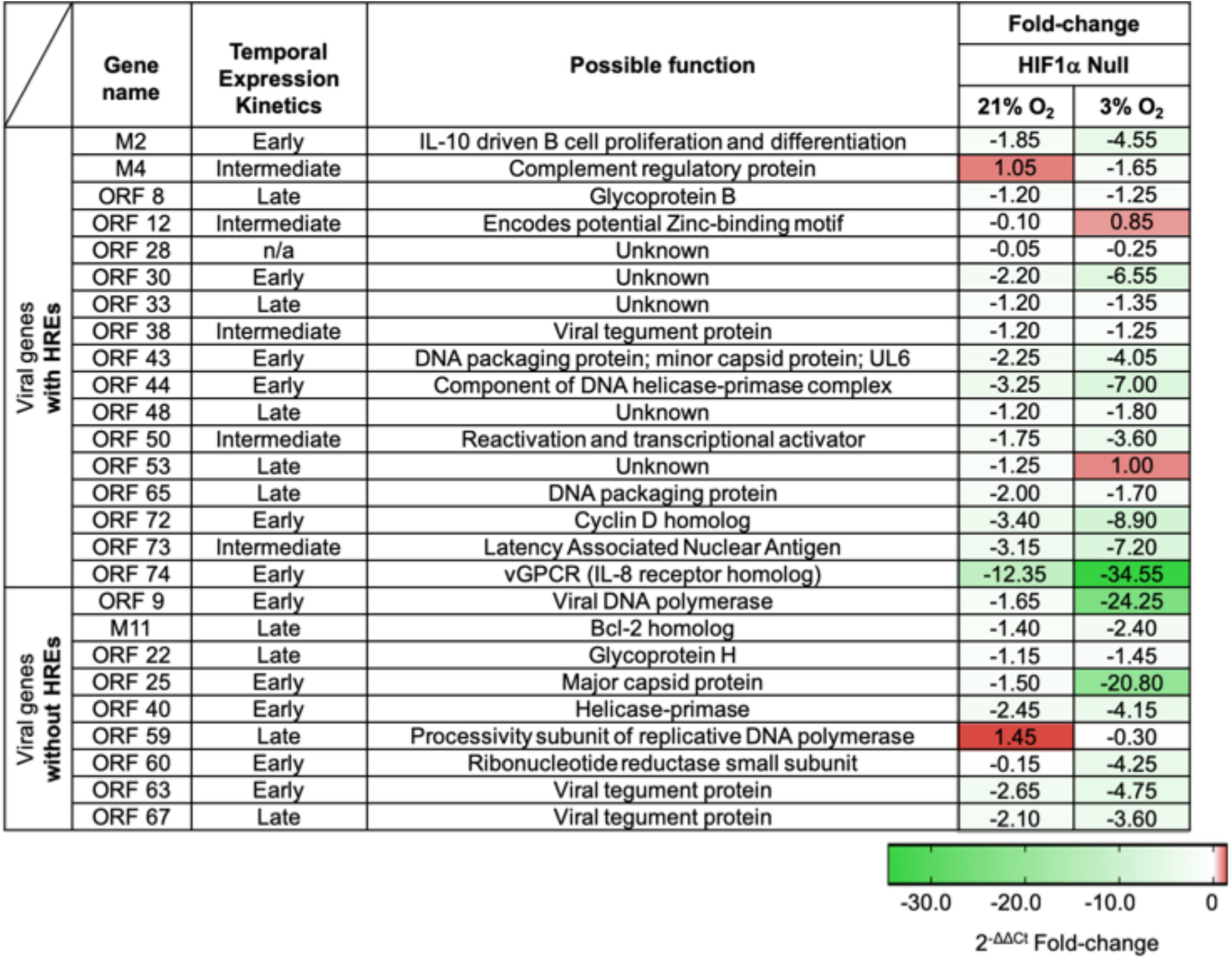
Gammaherpesvirus accelerates reactivation from latency and increased virus replication in physiological oxygen tensions. MHV68 latently infected splenocytes from C57bl/6J mice (n=3) were collected and processed on 16 dpi. Limiting dilution assay was performed on 21% O_2_ or 1% O_2_ to determine frequency of viral reactivation in low oxygen levels. At 63.2% the frequency reactivating splenocytes (dotted line) normoxia reactivation was 1 in 42,743 and from physioxic conditions 1 in 29,498. Symbols represent the mean percentage of wells positive for virus detection +/− standard error of the mean. Curve fit line were derived from nonlinear regression analysis. (**B-C**) Different ratios of splenocytes to 10^5^ MEFs, (10^4^ Left, 10^5^ Center, 10^6^ Right) were plated and incubated at 21% O_2_ or 1% O_2_. Graphs represent one out of two experiments. (**B**) Supernatants were collected, and viral titers were quantified by plaque assay on days 4, 5 and 6 post infection. *, *p<0.05. **, p<0.01. ****, p<0.001*. Statistical significance was determined by Multiple student’s t-test (**C**) At 3 dpi, RNA was isolated from cell layer of co-cultures and RTA levels were measured by qRT-PCR. *, *p<0.05*. Statistical significance determined by Student’s t-test.

## DISCUSSION

Understanding the role of the HIF1 pathway in the viral life cycle of oncogenic gammaherpesviruses is currently hindered by the lack of a suitable infection model. We present cumulative data indicating the importance of HIF1α in MHV68 lytic replication and reactivation from latency. In this study, we show that MHV68 activates the HIF1 pathway and that knock-out of *HIFIα* transcriptional activity diminished lytic replication *in vitro* and in an *in vivo* model of HIF knock-out of infected cells. Moreover, this truncated form of HIF1α impaired lytic reactivation of cells latently infected *in vivo*.

We show that MHV68 infection increased HIF1α protein levels. This was coupled with increase in HIF1α-dependent transcription activity (Fig. 1). A similar HIF upregulation was found in endothelial cells latently infected by KSHV [32], an oncogenic gammaherpesvirus that encodes many genes with the potential to upregulate HIF1α [13]. Although, the MHV68 viral genome is structurally similar to KSHV with many of the viral homologous [48] found to activate the HIF pathway in KSHV are also present in MHV68 but, the exact mechanisms whereby the virus could target the HIF1 pathway are still to be defined. A recent report shows that MHV68 activates IKKβ, a recently discovered transcriptional activator of HIF1α during cellular defense against microbes [49,50].

To further define the role of HIF1α in replication we generated knock-out cells (Fig. 2). We found that deletion of the DNA-binding motif impaired viral replication (Fig. 3A). In searching for possible explanations for the decrease in viral replication we analyzed the expression of viral HRE containing genes (Fig. 3B) and found that in 7 of 17 viral gene mRNA levels were decreased in the HIF deleted (Fig 3C). Our findings confirmed the conserved HREs within ORF50 previously published [27]. The majority of these HRE-containing viral genes have been described as essential for lytic replication such as DNA replication and assembly of mature virions [28] (Supplementary Table 1). Out of these genes, particularly striking was the downregulation of the KSHV homologue vGPCR gene in HIF1α Null Cells (Figure 3B). This is consistent with a recent report that also show that KSHV vGPCR is a gene under strong regulation by HIF1α [23]. Since vGPCR is required for RTA gene and protein expression during lytic replication of KSHV [51] it is possible that vGPCR downregulation in the absence of HIF1α, would hinder RTA participation in lytic gene expression.

We found that the upregulation of glycolytic enzymes by MHV68 during normoxic infection was impaired by HIF1α deletion (Fig. 3D). Metabolic reprogramming by gammaherpesviruses, was also observed during KSHV and EBV *de novo* infection *in vitro* [19]. Previous results published by our laboratory have demonstrated that treatment of MHV68 infected cells with 2-deoxy-D-glucose (2DG), a glycolytic inhibitor that was found inhibits the growth of KSHV-infected cells metabolically reprogrammed to glycolysis in normoxia [52], decreased by 5-fold of production of mature virions in comparison to untreated cells [53] pointing to a role of metabolic reprogramming involving HIF1α in MHV68 replication.

Recent studies have demonstrated that low oxygen tension can either enhance or downregulate virus infection [11,54]. It is unknown whether low oxygen level influences gammaherpesvirus lytic replication in the absence of HIF1α. We found that MHV68 lytic program is more heavily influenced by HIF1α under lower oxygen conditions—similar to physiological levels of oxygen in tissues and cells [55]. We found that at 3% oxygen levels, MHV68 in HIF1α Null cells undergoes a further significant replication impairment concomitant to a higher decrease on viral gene expression (Fig 4). Moreover, we found that expression of non-HRE viral genes were also significantly decreased during infection in Null cells (Suppl. Table 1). This could be the consequence of other transcription factors upregulated by an oxygen-depleted environment that could contribute to viral gene regulation [56,57] but; more likely the fact that many non-HRE containing viral genes are regulated by HIF-regulated genes such as RTA. Taken together our *in vitro* results in 21% O_2_ and 3% O_2_ show that HIF1α plays an important role in MHV68 replication and that this is due, at least in part, by a key role in the regulation of viral gene transcription.

Infection of floxed transgenic mice with MHV68-Cre to knock-out HIF1α *in vivo* revealed that HIF1α is necessary for optimal viral expansion on the site of acute infection in the animal model. This is consistent with our data of Figures 1 and 3, 4 showing that HIF1α upregulation plays a role in MHV-68 de lytic infection by regulating its lytic genes. This could explain why the peak of viral titers are decreased in lungs (Fig. 5B-5C). It is also likely that a reduction in viral expansion during the initial lytic phases in the lung could affect the extent of inflammation explaining the significant decrease of IL1β production in lungs lysates on day 7 (Figure 5D). These findings suggest that in the gammaherpesvirus life cycle, HIF1α is necessary for lytic virus expansion during acute infection of its host.

Similar to other herpesviruses, acute replication of gammaherpesviruses is followed by long-term establishment of latent reservoirs in the host. Although the frequency of latency establishment shown by nested PCR of bulk splenocytes on day 16 (Fig. 6A) was the same between wild type and HIF1αLoxP mice after MHV68-Cre virus infection, we found that the frequency of ex *vivo* reactivation of lytic virus from latently infected splenocytes was impaired in the absence of HIF1α (Fig. 6B). To further establish a possible role of HIF1α in reactivation from latency we tested WT virus reactivation in a lower oxygen context that we have shown increase HIF1α levels and activity. We found that low oxygen concentrations accelerated MHV68 reactivation and significantly increased the amount of infectious virions released concomitant with viral RTA upregulation. Our data further points to a critical role of HIF in gammaherpesvirus infection as it is likely to affect not only viral replication but viral reactivation in tissues where there are lower physiological oxygen level. It previously found, that exposure of KSHV, and EBV infected cells to hypoxic conditions can trigger a latent to lytic replication switch and enhance viral production and reactivation [11,58]. This likely happens via interaction of HIF1α with the transcriptional machinery that regulates viral expression. In KSHV, the expression of the Replication and Transcription Activation (RTA) and a lytic gene cluster is enhanced by HIF1α in complex with viral-encoded proteins such as the Latency-Associated Nuclear Antigen (LANA) [31,59]. Similarly, in EBV positive cell lines, HIF1α binds HREs located within the promoter region of the latent-lytic switch gene, *BZLF1* [12].

Both in KSHV and EBV, HIF1α has been linked to metabolic reprogramming of the host cell, modulation of viral latency, lytic replication and tumorigenesis. Our work further contributes to the understanding of HIF1 pathway during viral productive cycle in a natural infection and lytic replication in a cell and animal model. It establishes the utility of MHV68 as a model that can further our understanding the mechanisms whereby gammaherpesviruses interact with oxygen-sensing pathways. Our data also opens up new avenues to dissect the contribution of HIF1α in gammaherpesvirus infection of specific cell types such as myeloid and naïve and memory B cells that are targeted by MHV68 *in vivo* and can lead to lymphomagenesis under immunosuppression [60]. Our findings demonstrate the importance of the interplay of the oxygen sensing machinery and gammaherpesviruses, which is key to understand their pathobiology.

## METHODS

### Mice

Mice containing germ line floxed exon 2 of HIF1α gene (HIF1α^LoxP/LoxP^) on a B6.129 background were purchased from Jackson Labs and together with age- and sex-matched with wild-type C57BL/6 J mice were bred and maintained at our institute animal facility. Female mice at 8- to 12-week-old were used in groups of three to nine in most experiments. The animal experiments described here were performed according to the approved protocol by the University of Miami Miller School of Medicine Institutional Animal Care and Use Committee. In addition, we report compliance with the ARRIVE guidelines as requirement for reporting in vivo animal experiments.

### Virus stock

MHV68 containing Cre-recombinase (MHV68-Cre) driven by human cytomegalovirus promoter and parental MHV68-BAC virus were kindly provided by Dr. Samuel Speck, Emory University, Atlanta. MHV68-WUMS strain was obtained from Dr. Herbert Virgin, University of Washington, St. Louis. Viral stocks were prepared by low MOI infection of 3T12 cells in 2% FBS complete medium. Virus stocks were harvested after 5 to 7 days of infection and was processed by freeze-thawed cycle followed by homogenization. Subsequently, virus lysate was purified by centrifugation at 1,000 rpm for 10 minutes and supernatant was filtered thru 0.4μm membrane to remove cell debris. Finally, purified virus stock was prepared by ultracentrifugation at 27,000 rpm for 1 hour at 4°C and aliquots were transferred to −80°C for long-term storage. Viruses were quantified on 3T12 cells by standard plaque assay. Briefly, supernatants were diluted in 10-fold and transferred to cells layer in 24-well plates and incubated for 2 hours. 0.75% CMC containing overlay with 2% FBS complete media was added after inoculation. UV inactivation of viral stock was performed on a 60-mm plate in a Stratalinker, followed by plaque titration to ensure viral inactivation.

### Cell culture

NIH 3T12 (ATCC CCL-164), a fibroblast cell line permissive to MHV68 replication was used to test the status of HIF1α after infection with MHV68-WUMS Strain. For all subsequent in vitro studies to test the absence of HIF1α activity, murine embryonic fibroblast (MEFs) were immortalized using the 3T3 NIH method. Briefly, MEFs were obtained from C57BL6 and B6. HIF1α^2loxp^ at 13.5 to 15.5 days post-coitus and cultured in T-25 flasks at 3X10^5^ cells every 3 days until passage 32. Immortalized HIF1α^LoxP/LoxP^ MEFs were generated by lentiviral transduction of Cre-recombinase and selected by Blasticidin. MEFs and 3T12 were cultured at 37°C with 5%CO_2_ in Dulbecco modified Eagle medium (DMEM) containing 10% fetal bovine serum (FBS), 2μM L-glutamine, 10μg/ml of gentamicin.

### Low oxygen treatment

In experiments that required low oxygen concentration, cells were cultured in a humid hypoxia chamber under a mixture of O_2_/ CO_2_/ N_2_. To obtain physiological oxygen concentrations or 3% O_2_ conditions, 3:5:92 vol% and 1% O_2_, 1:5:94 vol%. Hypoxia mimic was achieved by treatment with Cobalt chloride (Roche) at 150μM.

### Excision assay

Cells were lysed in RLT buffer (Qiagen) supplemented with 1% β-mercaptoethanol and stored at −80°C before RNA extraction. RNA was isolated using RNeasy minikit (Qiagen) and cDNA was prepared using ImProm-II Reverse Transcription System (Promega) according to manufacturer’s instruction. PCR conditions were as follow 95°C for 2 minutes followed by 32 cycles of 95 °C for 30 seconds, 64°C for 45 seconds and 72°C for 45 seconds. Excision was demonstrated by a shift in the size the mRNA fragment spanning exon 1 to exon 5 (600bp), which upon deletion of exon 2 can be detected in a 2.5 % DNA agarose gel as a 400bp fragment when amplified by PCR (Invitrogen). Wild type MEFs isolated from C57Bl/6J mice transduced with the cre-recombinase expressing lentivirus was used as control to detect the specificity of excision in floxed HIF MEFs. Primer sets were purchased from Sigma at follows: exon 1 forward 5’-CCGGCGGCGAGAAG −3’ and exon 5 reverse 5’-CCACGTT GCT GACTT GAT GTT CAT- 3’.

### Western blot

Samples were lysed in RIPA buffer and sonicated to avoid clumps from genomic DNA in lysates. Protein concentration was determined with BCA assay (Thermo Scientific) prior to resuspending in Laemmli buffer. 20 ug of protein lysates were separated by SDS-PAGE. Proteins were transferred to a PDVF membrane (Pall Life Sciences) at 100V for 1 hour at 4°C. Membrane was probed with 1:500 of mouse HIF1α antibody (Novus biologics) and 1: 2,000 mouse beta-actin antibody (Sigma). Primary antibodies were detected with HRP-conjugated anti-mouse secondary antibody (Sigma) and chemiluminescence reagent (Thermo Scientific).

### Reporter assay

Transfection of 3T12 and MEFs were performed by using Lipofectamine 2000 following manufacturer’s protocol. The HRE-Luciferase reporter is a pGL2 vector containing three hypoxia response elements from the *Pgk-1* gene upstream of firefly luciferase [61]. TK-Renilla (0.5 μg per well) and HRE-luciferase (0.5 μg per well) was co-transfected to control for transfection efficiency. After 12 hours of transfection, cells were infected with MHV68 at low MOI of (0.5 PFU/cell) and high MOI (3.0 PFU/cell). Firefly luciferase (F-Luc) activity in cell lysate was measured using Dual Luciferase Assay System (Promega Corporation), as recommended by the manufacturer. Renilla luciferase (R-Luc) activity in cell lysates was measured using 12 M coelenterazine in assay buffer (50 mM potassium phosphate, pH 7.4, 500 mM NaCl, 1 mM using a luminometer. Data is expressed in relative light units and percentage of HIF1α dependent transcription was determined by normalizing luciferase units to Renilla.

### Real-Time qPCR

Cells were lysed in RLT buffer (Qiagen) supplemented with 1% β-mercaptoethanol and stored at −80°C before RNA extraction. RNA was isolated using RNeasy minikit (Qiagen) and cDNA was prepared using ImProm-II Reverse Transcription System (Promega) according to manufacturer’s instruction. Quantitative PCR was performed with 10 to 50 ng of cDNA using SyBr Green (Quanta Biosciences). PCR conditions were 95°C for 5 minutes followed by 45 cycles of 95 °C for 10 seconds, 60°C for 20 seconds and 72°C for 30 seconds. The TATA-binding site mRNA was used as housekeeping gene. We compared the normalized Ct values (ΔCt) of each gene in two biological replicates between two groups of samples. All relative fold-change values were normalized against normoxic conditions using 2^-ΔΔCt^ to display fold-change.

### *In vitro* viral infections

Viral infections were performed in low volume serum-free complete media at 4°C for 2 hours at 21% O_2_. Cell layers were washed twice with 1X PBS and then 2% FBS complete medium was added for experiments. For RNA and protein analysis, cell layer was washed once with cold 1X PBS.

### Viral pathogenesis assays

Wild type and HIF1α-LoxP mice were euthanized with ketamine (100mg/kg) and xylazine (10mg/kg) and infected with 3×10^4^ PFU of MHV68-Cre virus. Lungs were removed on days 3, 5 and 7 post infection and freezed-thawed prior to processing. Tissue was disrupted in 1ml of 2% FBS complete medium using a handheld Omni homogenizer (www.OMNI-INC.com). Viral titers were determined by plaque assay on 3T12 cells plated in 24 well-plates and cultured in 0.75% CMC-overlay medium.

### Limiting dilution assay

Bulk splenocytes were serially diluted by 2-fold on days 16 and 42 post infection after RBC lysis and plated on primary MEF starting at 1X 10^5^ cells/well down to 7.5X10^2^ cells/well with replicates of 24-well per dilution in 96-well plate. After 3 weeks, sups were collected and re-plated into 3T12 cells to amplify the virus and cytopathic effects was scored.

### Viral genome frequency

Splenocytes were thawed and counted and diluted in 10^4^ uninfected 3T12 cells. After proteinase K treatment, two rounds of PCR were performed against MHV68 ORF50. Copies of a plasmid containing ORF50 in 10, 1 and 0.1 copies were diluted against 1X10^4^ 3T12 cells and amplified in each run to ensure sensitivity of assay.

### Virus reactivation of splenocytes in low oxygen

At day 16 following intranasal infection, (n=3) spleens were processed to obtain splenocytes at a single-cell suspension. Explanted splenocytes were plated at different quantities (10^4^, 10^5^ and 10^6^) on top of a MEFs layer (1X10^5^ cells per well) in a 6-well plate in duplicates. The-co-culture was kept in 2-ml of 2% FBS complete 1X DMEM media then transfer to 21% O_2_ or 1% O_2_ conditions.

### Identification of HRE sequences within viral sequences

Computer-assisted prediction of HIF1α binding sites within the 500bp upstream of MHV68 ORFs was performed with TESS (Transcription Element Search System) using TRANSFAC for the search string RCGCT allowing only core position for strings with a maximum allowable string mismatch of %10.

### Enzyme-linked Immunosorbent Assay

IL-1 β and TNF-a were quantified by a mouse ELISA Ready-SET-Go! Kit (Affimetrix, eBioscience San Diego, CA). Plates were prepared and assayed according to the manufacturer’s protocol and signals were read at 450 nm and subtracted the values of 570 nm to those of 450 nm.

### Statistical analyses

Data analysis was perform using Prism software (Graphpad).

Viral titer, reporter assays and mRNA fold-change was analyzed with a two-tailed Student *t* test and values are expressed as the means of standard error. Frequencies of reactivation and viral DNA positive cells were determined within the nonlinear regression fit of the results on the regression line that intersected at 63.2%, following a Poisson distribution. Results were considered to be statistically significant for values of P<0.05.

## Acknowledgements

Funding for this work was provided by through a development award from the National Institute of Allergy and Infectious Diseases, Center for AIDS Research, University of Miami, P30A1073961 to SA and EAM and by NIH grant CA137386 to EAM. We are grateful to Dr. Samuel Speck and Dr. Herbert Virgin for providing the MHV68 viruses. We acknowledge Dr. Clinton Padden for his technical advice on *in vivo* latency experiments and David Wilde and Adriana Correa for their assistance in performing virus titers and RNA extraction. The HRE-analysis of viral genes was performed in collaboration with Dr. Daria Salyakina of the Oncogenomics Core at the University of Miami. We would like to thank Julián Naipauer, Santas Rosario, Omayra Méndez and Frances Collins for their feedback on the manuscript.

## Author Contributions

### Conceptualization

DMLR EAM SA

### Formal analysis

DMLR LTK EAM SA

### Funding acquisition

EAM SA

### Investigation

DMLR VK LTK EAM SA

### Methodology

DMLR VK LTK EAM SA

### Project administration

EAM SA

### Resources

EAM SA

### Supervision

EAM SA

### Writing- original draft

DMLR LTK EAM SA

### Writing- review and editing

DMLR EAM SA

---

**Supplementary Table 1. Absence of HIF1α impairs gammaherpesvirus gene in low oxygen levels.** HIF1α WT and HIF1α Null MEFs were infected with MHV68 (MOI 5.0) and transferred to either 21 % and 3% oxygen, RNA was isolated 24 hpi. Levels of mRNA for MHV68 ORFs with HRE were determined by qPCR; ΔΔCt normalized against WT infection at 21% O_2_ and displayed as 2^_ΔΔCt^ fold-change. Heat map was created using GraphPad Prism. Statistical significance displayed as asterisk (*, *p<0.05*) were determined using GraphPad Prism by Bonferroni’s multiple-comparison test as following: column 1: 21% O_2_ HIF1α Null vs 21% O_2_ HIF1α WT, column 2: 3% O_2_ HIF1α Null vs 21% O_2_ HIF1α WT.

